# Small-scale biogeography of biofilms and its implications for sequencing-based studies

**DOI:** 10.1101/2025.10.13.682146

**Authors:** Marco Gabrielli, Frederik Hammes

**Author notes:** Corresponding author: Name: Marco Gabrielli.

## Abstract

Biofilm microbial communities are characterized by spatial heterogeneity arising in response to environmental characteristics and the metabolic requirements of their inhabitants, which create biogeographical patterns at both large and small scales. While the impact of small-scale biogeography on several emergent biofilm properties has been characterized by a variety of laboratory studies using simple synthetic communities, the extent of small-scale biogeography in environmental biofilms, as well as its impact on sequencing-based studies, remain poorly characterized. Here, we investigate centimeter-scale biogeography in potable water biofilms developed in environments with distinct levels of heterogeneity. We then estimate microbial diversity at scales relevant for microbial interactions and characterize the impact of sampling strategies on both alpha and beta diversity metrics, as well as on inferred co-occurrences and scaling laws. Our results show that biofilm sections in seemingly homogeneous environments are characterized by similar biogeographic patterns as the ones in a more heterogeneous one, albeit with lower cm-scale variability. We estimated that, despite the overall richness over the entire biofilms, single cells are surrounded by local neighborhoods of limited diversity, strongly limiting their possibility for diverse interactions. Larger sampling areas average such local heterogeneities and distort community structure estimates, leading to an inflated and erroneous estimation of co-occurrences and potential metabolic interactions. Finally, we report how both environmental heterogeneity, as well as sample area size, influence scaling law performances. Our results highlight the ubiquitous nature of biofilm biogeography and call for the adaptation of biofilm sampling strategies to consider this phenomenon relative to specific research questions.

## Introduction

Biofilms contain the majority of biomass on Earth and are responsible for key ecosystem services [1]. The microbial communities residing in biofilms are affected by the environmental conditions in their surroundings, resulting in location-specific composition which generate regional and global biogeographic patterns [2,3]. Besides these large-scale differences, small-scale differences develop within biofilms due to the geometric and physico-chemical properties of their environments [4], as well as the presence of chemical gradients of nutrients and other specialized molecules arising from the metabolic activity of their inhabitants [5,6]. Such local heterogeneous conditions can produce different spatial configurations in the microbial communities, which influence emergent properties of biofilms such as virulence and resistance to treatment [7], as well as horizontal gene transfer and metabolite production and consumption [8–10]. While the causes of these spatial configurations are relatively simple (i.e., growth and movement), biogeographic patterns in complex communities are still largely unexplored due to the prevalence of laboratory studies involving a very limited number of species and the complex interdependencies of their causes (e.g., interactions types, fluid flow, cell type) [11].

Studies focusing on the small-scale biogeography of environmental biofilms usually rely on FISH-based approaches [12,13]. While this technique allows to obtain a precise understanding of the spatial configurations of the microbes of interests, it can only track a limited number of different taxa compared to the diversity of environmental biofilms, providing limited community-level insights [14]. In contrast, sequencing-based studies have largely ignored the presence of small-scale biogeography [15] and focus on differences across large environments such as watersheds [3]. However, in case of environments with high degrees of small-scale heterogeneity (e.g., soil) the relevance of large sample areas/volumes to study microbial interactions and ecosystem services is questionable [16]. Besides such heterogeneous environments, even ones assumed to be relatively homogeneous like potable water distribution systems have been shown to harbor heterogeneous biofilms [17,18], suggesting that, regardless of the type of environment, spatial patterns are a key feature of biofilms. Hence, given the pervasiveness and importance of biofilms, as well as the widespread use of sequencing approaches for microbiology research, it is of primary importance to understand how sampling strategies affect our microbiological understanding of microbial biofilms and their small-scale biogeography.

The goal of this study was to characterize the small-scale biogeography of biofilms in environments with distinct levels of environmental variability and evaluate the impact of sampling area on the estimated microbiological characteristics of biofilm communities. To do this we re-analyzed and explored the dataset from Neu et al. [18] where two biofilms developed in environments with different levels of environmental heterogeneity were sequenced at cm-scale resolution to characterize the degree of heterogeneity in biofilm composition. Then, we used the same data to estimate the diversity of microbial neighborhoods at micrometer scale and test the variations of diversity metrics, inferred interactions and scaling laws as a function of sample area.

## Materials and methods

### Datasets used

The dataset of Neu et al. [18] was derived from the biofilm of two shower hoses made of plasticized polyvinyl chloride (PVC-P), with an inner diameter of 0.8 cm and a total length of 1.80 m. One hose was kept horizontal and undisturbed to prevent physical biofilm disruption, and flushed regularly with warm water (35–42°C) twice per day, minimizing as much as possible external disturbances. The other hose was installed in a volunteer house and was subjected to normal usage, kept mostly vertical and subjected to variable flow velocities and water temperatures, movement, uncontrolled stagnation and potentially partial drainage. Upon collection, both hoses were cut in pieces bisecting the pipe in length (i.e., top/bottom and side1/side2) and with individual lengths of 1.2 cm splitting the hoses in 200 biofilm sections. For each biofilm section, the cell number was measured with flow cytometry, while the prokaryotic community composition was assessed using 16S amplicon sequencing of the V3-V5 region (for complete details, see: Neu et al., [18]).

### Amplicon sequencing analysis pipeline

Downloaded raw reads were subjected to quality control by checking for the presence of PhiX contamination, removing low complexity reads and low-quality read lengths using USEARCH v11.0.667 [19] and removing primers with Cutadapt v4.9 [20]. Reads were merged into contigs using USEARCH, filtering the ones with more than one expected error. Finally, zOTUs were retrieved thanks to the UNOISE3 algorithm [21], as implemented in USEARCH. zOTUs taxonomy was estimated using SINTAX [22] based on RDP database (release No. 19) [23], while phylogeny was estimated using IQTREE v2.2.2.6 [24] using a GTR substitution model on an alignment of the zOTUs obtained thanks to MAFFT L-INS-i v7.505 [25] and TrimAl v1.4.rev15 [26]. Long branching zOTUs were manually inspected and eventually excluded from the datasets. Contaminant zOTUs were removed exploiting the negative control samples sequenced with the package decontam v1.22.0 [27]. Finally, samples were rarefied to an even sequencing depth through the phyloseq v1.46.0 [28] package in R v4.3.0 [29]. Highly related zOTUs were clustered in OTUs at 99% percentage identity using DECIPHER v2.30.0 [30].

### Microbial community characterization

For each biofilm pieces and pair of pieces belonging to the same dataset, alpha diversity metrics (i.e., observed richness, Shannon index, Simpson index, Pileou evenness) and beta diversity (i.e., Bray-Curtis dissimilarities) were estimated using the packages phyloseq, microbiome v1.24.0 [31] and vegan v2.6-4 [32]. Beta diversity was estimated based on both the total microbial community, as well as separating its core and satellite members, identified based on their abundance-occupancy distributions as proposed by Shade and Stopnisek [33]. zOTUs with monotonic relative abundance or non-random trends were identified using, respectively, Spearman correlations and Wald–Wolfowitz runs test correcting the obtained p-values through the Benjamini-Holchberg method. Functional traits (i.e., KEGG orthologs, KO) were inferred using PICRUSt2 v2.6.0 [34] relying on the GTDB r214 database [35]. Differences in correlation coefficients were tested thanks to the ad-hoc statistical tests implemented in the R package cocor v1.1-4 [36]. Changes in the patterns of beta diversity along both environments were detected using the package segmented v2.0-3 [37].

### Estimation of diversity in local microbial neighborhoods

Total cells concentrations and biofilm thickness measurements presented in Neu et al. [18] were used to estimate the expected number of cells within neighborhoods of different areas or volumes. Such values, together with the zOTUs relative abundances in the respective biofilm section, were used to estimate the number of different zOTUs expected within such neighborhoods leveraging the Heaps’ Law solution proposed by Ferrante and Frigo [38] (Eq 1), where Ε[𝑅_𝑚_(𝑛)] is the expected richness in a given neighborhood, 𝑚 is the total richness in a biofilm section, 𝑛 is the number of cells in a given neighborhood and 𝑝_𝑖_ is the relative abundance of zOTU 𝑖 in a biofilm section (i.e., the probability of selecting it randomly).

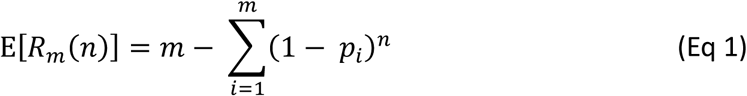

### Evaluation of alternative sampling strategies

Contiguous or randomly-selected biofilm sections were merged to simulate alternative sampling strategies with different sample areas. The newly generated samples were rarefied to the same sequencing depth as the original samples, mimicking a scenario where either a commercial sequencing platform with a fixed sequencing depth per sample is chosen or one in which the fewer number of samples per specific environment would be counterbalanced by a larger number of samples across environments (e.g., total sequencing depth for a study capped). For each simulated sampling strategy, alpha diversity metrices was estimated both for each simulated sample, as well as the entire hose. Similarly, beta diversity among simulated samples was also estimated. Finally, the number of zOTUs present in all samples, as well as the number of statistically co-occurring zOTU pairs was obtained thanks to the package CooccurrenceAffinity v1.0.2 [39]. To control the effect of a different number of samples on the co-occurrence estimates, all sampling strategies were tested by selecting randomly an equal number of samples and iterating this process 100 times.

### Scaling laws estimation

Richness scaling laws were fitted based on the power law (*RPL*) and the logarithmic power laws (*RLPL*) (Eqs. 2) where *A* represent the biofilm area, *z* the estimated spatial turnover rate, while *b* and *c* other shape parameters [40]. This was performed on both the real samples from both hoses, as well as the simulated samples with larger area fractions. Accuracy of the scaling laws was measured through the root mean square error (RMSE) against the species-area relationship (SAR) curves estimated through the package vegan, accounting for the different sample area among the simulated samples. Due to the variability of SAR curves caused by sample randomization, RMSE was estimated 100 different estimates of such curves.

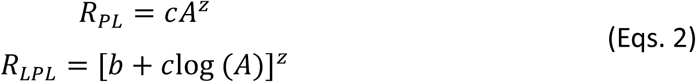

To provide statistical support, microbiome data emulating both biofilms was simulated 100 times using MIDASim v0.1.0 [41] in its nonparametric mode.

## Results

### Biofilms in controlled environments show comparable but less noisy biogeographic patterns compared to uncontrolled environments

In order to study the degree of biogeography as a function of environmental heterogeneity, we compared the community characteristics along two biofilms derived from environments with different degrees of heterogeneity. One biofilm was obtained from a shower hose that was kept undisturbed for a year. Given the lack of external disturbances and macroscopic homogeneity, it will be referred hereafter as the “controlled environment”. The second shower hose was subjected to normal but uncontrolled usage which, together with the vertical orientation of the hose, is likely to have resulted in temperature and humidity gradients along the hose. As a result, this hose presents heterogeneous environmental conditions and will be hereafter referred to as the “uncontrolled environment”. Differences in environmental heterogeneity resulted in different variabilities of biofilm communities along the two hoses (Figure 1), with a drastically more homogeneous core microbial community (i.e., the taxa presenting high relative abundance and occupancy) (Figure S1) in the controlled environment than in the uncontrolled one. Both environments showed taxa with relative abundance patterns which consistently increased or decreased along the environments’ length or peaked at specific locations (Table S1). In the uncontrolled environment, we detected these spatial patterns for a total of 148 of the 558 zOTUs (26%). Even though the controlled environment can be considered macroscopically homogeneous, we observed spatial patterns for 99 of the 502 zOTUs (20%) present. Approximately 83% of the zOTUs belonging to the core microbial community in the controlled environment (Figure 1A), as well as all of the 22 zOTUs in the uncontrolled one (Figure 1B) presented spatial patterns, indicating that these patterns are not caused by rare taxa. High upstream relative abundance was linked to both increases and decreases of relative abundance along the biofilm (Figure S2), suggesting that biofilm self-seeding did not play a major role in the final composition of biofilm. Finally, we observed that the co-occurrence of highly related zOTUs (i.e., clustered in 99% similarity OTU) in the same biofilm section occurred systematically (i.e., at least in 50% cases) for less than 30% of the zOTUs showing more than 99% similarity (Figure S3).

**Figure 1.**
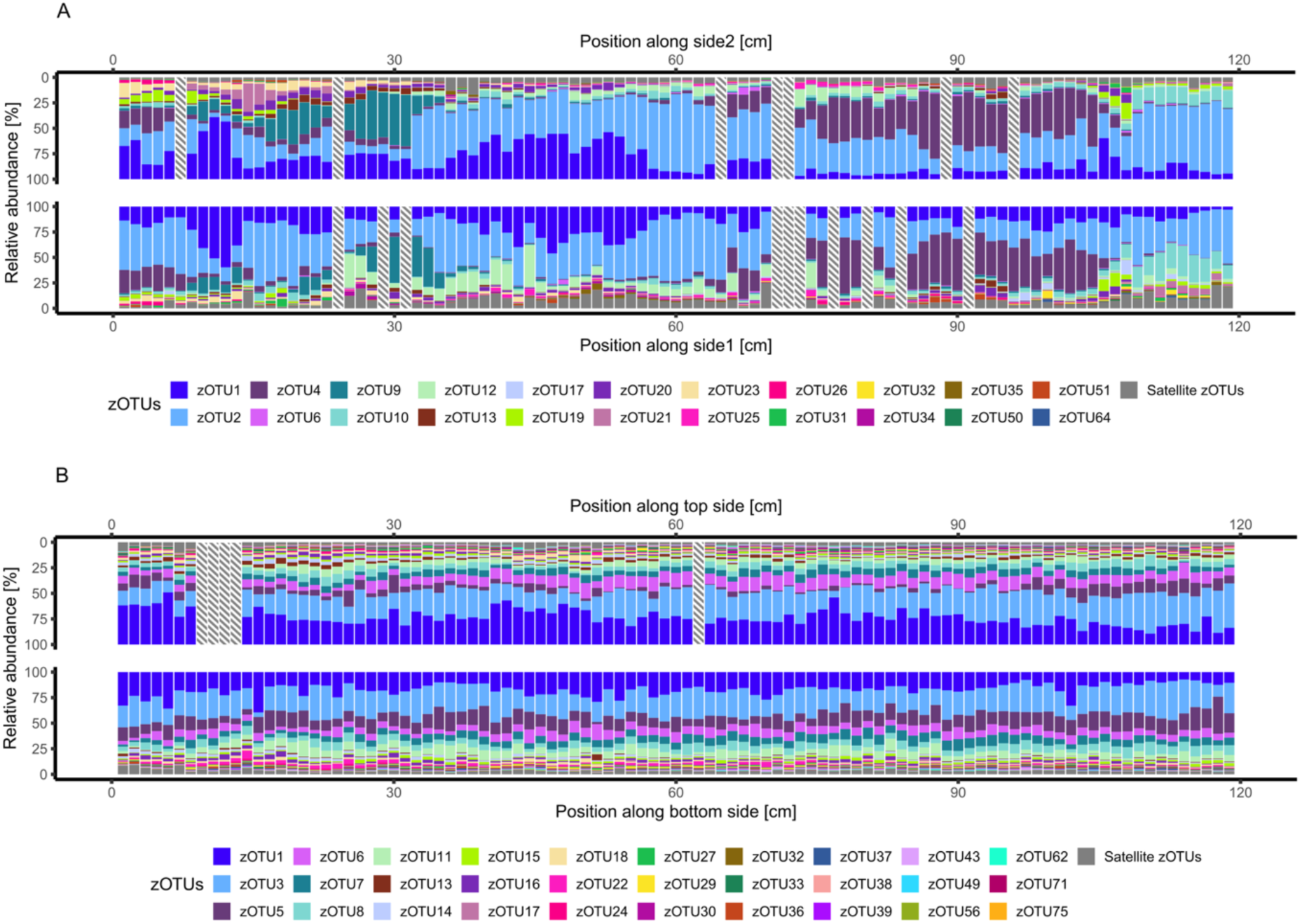
Biofilm microbial community structure on cm-scale in the uncontrolled environment with heterogeneous environmental conditions (A) and the controlled one macroscopically homogeneous conditions (B). Each sample represents a 1.2 cm biofilm section retrieved from the shower hoses bisected into two sides (i.e., side1/side2, top/bottom). Taxa not included in the core microbial communities of either environment were grouped as “Satellite zOTUs”. Striped sections indicate missing data.

The relative abundance patterns reflected themselves on the community composition along both biofilms. Regardless of the environment, the community structure changed along the length of the biofilms, showing significant correlations between alpha diversity metrics and position within the biofilm (Figure 2A,B; Figure S4 Table S2). The extent of the correlations was predominantly unaffected by environment heterogeneity, with the majority (62.5%) of the correlations between alpha diversity metrics and position along the biofilms being comparable between the two environments (Fisher test; p-val > 0.05). Yet, we found the alpha diversity in the uncontrolled environment showed greater variability between consecutive biofilm pieces (Wilcoxon test; p-vals < 0.003). In addition, we found that the orientation of the environment significantly affected community structure (Wilcoxon test; p-vals < 0.001). Given the horizontal orientation the controlled environment, this variation could be attributed to particle settling which would concentrate taxa and micro-nutrients associated with suspended solids to the bottom. While no explicit explanation could be found for this difference in the uncontrolled environment, the effect on the observed richness was significantly smaller than in case of the controlled environment (T-test, p-val < 0.001), suggesting the presence of only minor differences in conditions potentially caused by distinct shear stresses between the two sides of the hose.

**Figure 2.**
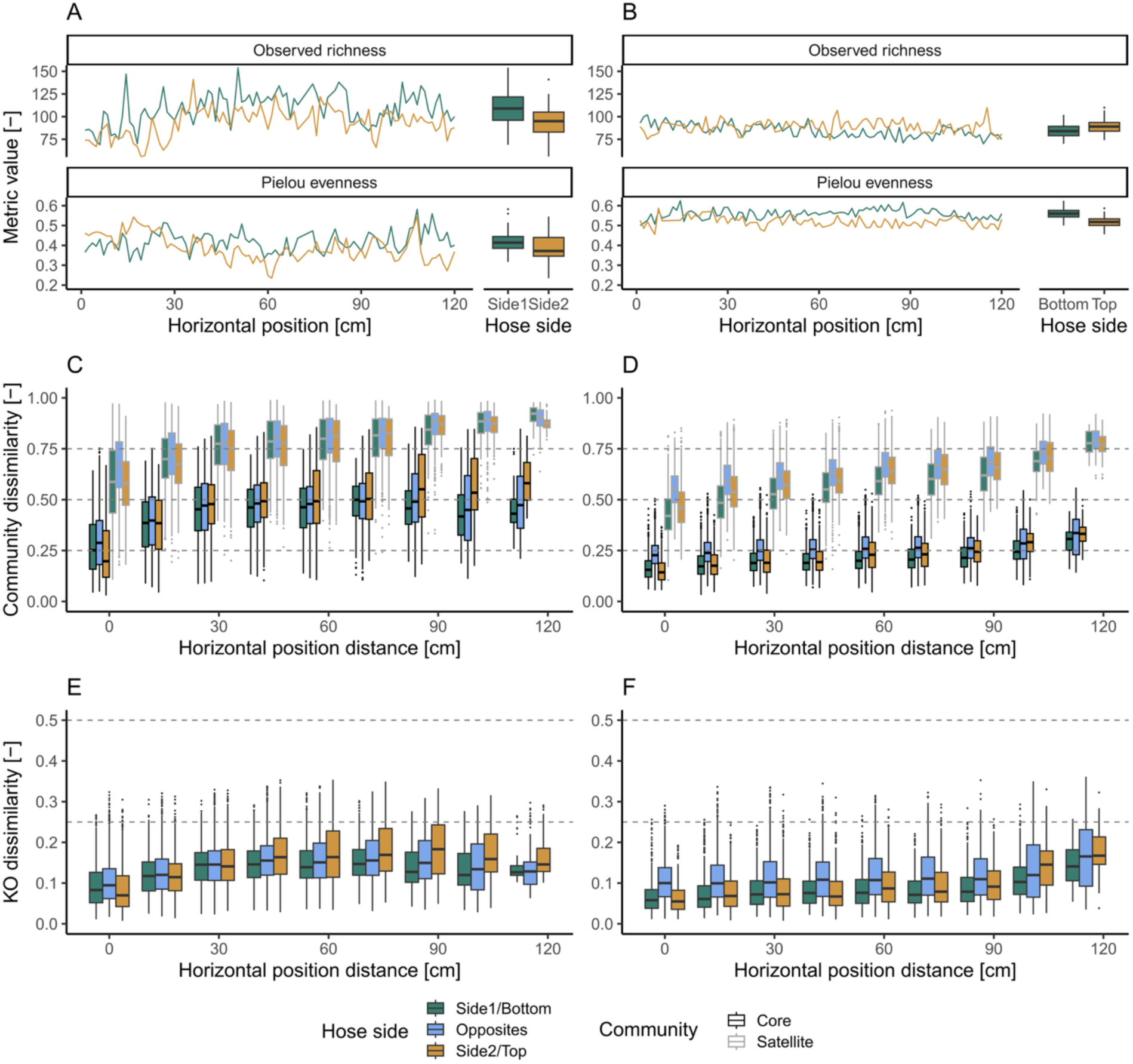
Alpha diversity metrices in the uncontrolled (A) and controlled (B) environments split by shower hose side along the shower hoses length. Bray-curtis dissimilarity of the core and satellite microbial communities in the uncontrolled (C) and controlled (D) environments as a function of distance between biofilm sections along the shower hoses. Bray-curtis dissimilarity of predicted KO profiles in the uncontrolled (E) and controlled (F) environments as a function of distance between biofilm sections along the shower hoses.

The composition of microbial communities changed within both environments, with closer sections of biofilm showing more similar core and satellite communities than distant ones (Figure 2C,D). The macroscopic homogeneity of the conditions in the controlled environment did not eliminate biogeographic patterns (i.e., the spatial distribution of taxa in an environment), but only reduced them. We found significantly lower beta diversity between biofilm sections at the same distance (ANOVA – Tukey HSD posthoc test; p-val < 0.001), as well as lower variability (ANOVA – Tukey HSD posthoc test; p-val < 0.001) in the controlled environment compared to the uncontrolled one. Notably, the difference among the beta diversity of the core and satellite community and the distance of biofilm sections did not increase at greater distances (Spearman correlation; p-val > 0.51), indicating comparable rates of change of the relative abundances of both common and rare microorganisms. Similarly to what noted for the alpha diversity, the different orientation of the sides in the controlled environment led to greater beta diversity than biofilm sections on the same side and at the same horizontal distance (ANOVA, p-val < 0.001), a pattern not observable in the uncontrolled environment.

Finally, we estimated the spatial differences of the inferred profiles of KEGG Orthology (KO) identifiers representing the inferred metabolic capabilities of the microbial communities in each section of the biofilms. While a direct comparison of the dissimilarities of KO profiles and community composition is not immediately possible due to the different nature of the two classifications, we assessed the distance at which the spatial turnover (i.e., the slope between dissimilarity and distance) changes significantly in both environments. Such changes occurred at comparable distances in the controlled environment (zOTUs: 92.4 ± 1.7 cm, KOs: 92.0 ± 1.3 cm) due to an increased spatial turnover at relatively large distances. On the other hand, in the uncontrolled environment, the KO profiles diversity reached a plateau at a significantly farther distance than microbial community membership (zOTUs: 22.3 ± 0.5 cm, KOs: 37.7 ± 1.0 cm), indicating that community functions are more spatially conserved than taxonomy. Yet, changes in the diversity patterns were found at lower distances than in the controlled environment, once again highlighting the effect of heterogeneous environmental conditions on microbial community membership and its encoded functions.

### Local neighborhoods show limited diversity

Contrary to most laboratory biofilm models, where the volume is mostly composed of cells, most of the volume of environmental biofilms can be composed of EPSs and inorganic material [42]. Accordingly, we found between 0.7 and 3.87×10^5^ cells within areas of 10 µm^2^ and 1 mm^2^ for both biofilms (Figure 3A), confirming the relative low cell density reported in the original publication [18]. As a result of such low cell densities, we estimated that maximum 3 different taxa to be present in a 10 µm^2^ area, a value concordant with biofilm imaging results (Figure 3E). On the other hand, a number of taxa compared to full biofilm sections (Figure 2) was predicted for 1 mm^2^ areas. Such values are likely overestimation of the real diversity in such areas given that Heaps’ Law does not take into consideration that prokaryotes reproduce by binary fission which results in daughter cells being co-localized with their parent cell (as observable in Figure 3E by the presence of cells of similar shape in proximity one to another).

**Figure 3.**
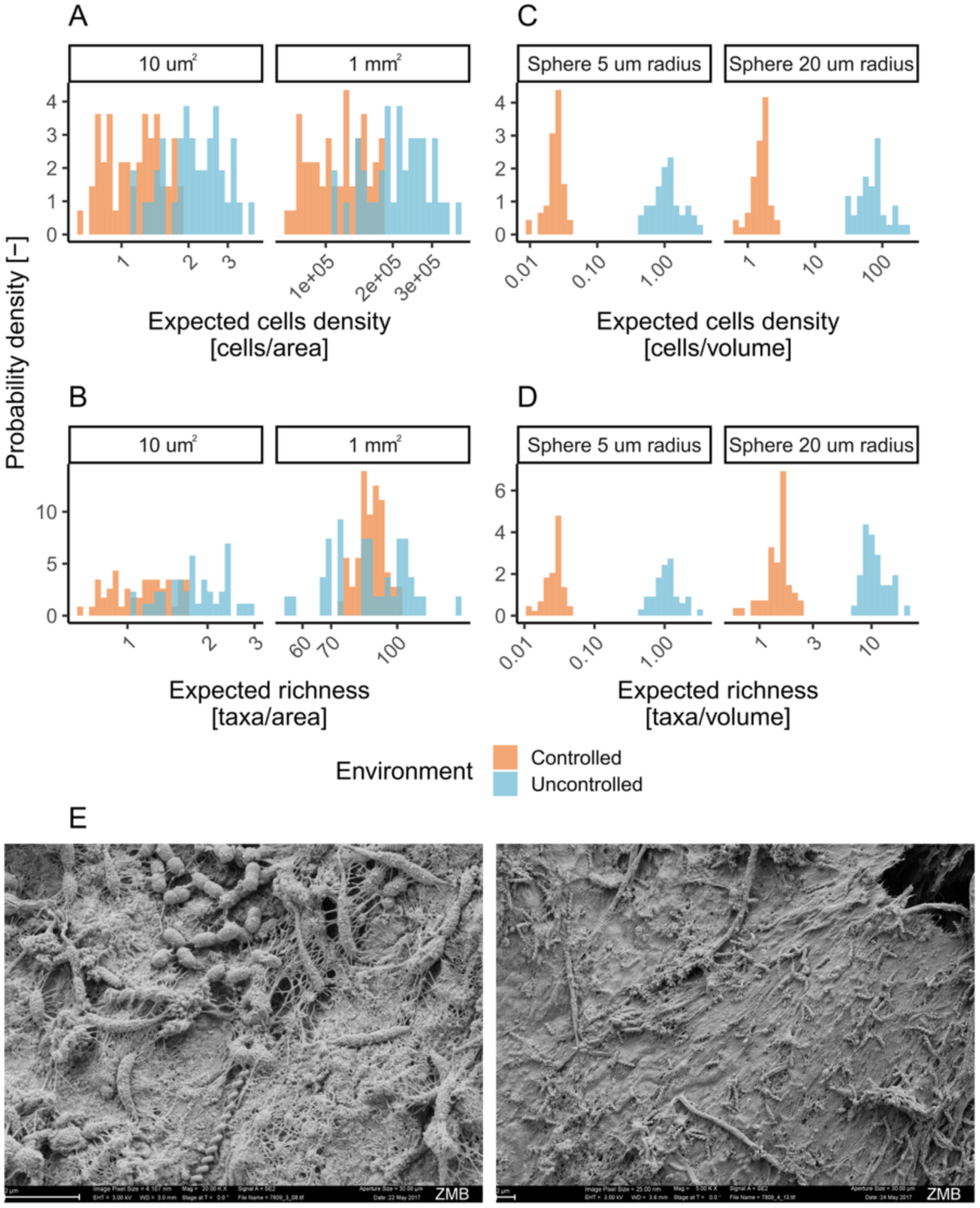
Number of cells expected in biofilm areas of 10 µm^2^ and 1 mm^2^ (A) and biofilm volumes within 5 and 20 µm radii (C). Estimated taxonomic richness present in such areas and volumes (B, D). Scanning electron micrographs from the biofilm developed under controlled conditions (E).

We estimated the number of cells and taxa around a single cell within tridimensional neighborhoods of radii between 5 and 20 µm, i.e., the range of distances compatible with metabolic exchanges [43]. Such analysis revealed that less than 3 and 22 different taxa were expected to be found, respectively in neighborhoods with a 5 and 20 µm radius (Figure 3C,D) with more local diversity in the uncontrolled environment. This is likely due to the fact that more turbulent flow, as expected in the uncontrolled environment, lead to denser biofilms due to the effect of shear stresses [44]. In any case, the low numbers of both cells and taxa found in both environment types indicate that the opportunities for contact-dependent or metabolic interactions of each single cell are rather limited. Yet, it is important to note that our estimates capture the diversity of microbial neighborhoods experienced by a single cell and not whole taxa. Assuming a random deposition on biofilms, two cells belonging to the same taxa are likely to experience different microbial neighborhoods, enabling greater possibilities for interaction among taxa.

### Trade-offs between specificity and representativeness in biofilm sampling

Given that our result shown that biogeographic patterns and spatial heterogeneities are inevitable within biofilms, it is evident that the sampling strategy will affect the microbiological observations of an environment. To test this, we simulated for each environment type two different sampling strategies with increasing biofilm area per sample and compared their results with the actual sampling strategy from the original study. One strategy, referred to as “contiguous” simulated sampling entire contiguous hose sections. Conversely, the strategy named “discontiguous” simulated a scenario in which randomly selected hose sections are pooled together in a single sample with the goal to reduce the influence of biofilm biogeography on the observed results.

Regardless of the environment characteristics or the sampling strategy, sequencing larger areas lead to an increase in all alpha diversity metrics per sample (Spearman’s correlation; rho > 0.15, p-values < 0.001), showing, however, only negligible effect on the value of Pielou’s evenness (|rho| < 0.09, p-values > 0.03) (Figure 4A). Coincidentally, we detected a decrease in the variability of all alpha diversity metrics when the sample area was increased (Spearman’s correlation; rho < -0.89, p-values < 0.001), except for the observed richness. Increasing sample area while fixing the sequencing depth led to a significant decrease in the observed richness over both environments, moving from approximately 500 zOTUs observed across both biofilms using the sampling scheme of the original publication to around 100 zOTUs when sampling the whole environment in a single sample. Such a decrease is caused by the loss of low-abundant taxa, as highlighted by the limited difference in Shannon and Simpon diversity and the drastic change in Pielou’s evenness before and after rarefaction. The selection of random samples throughout an environment simulated in the discontiguous strategy did not impact greatly alpha diversity.

**Figure 4.**
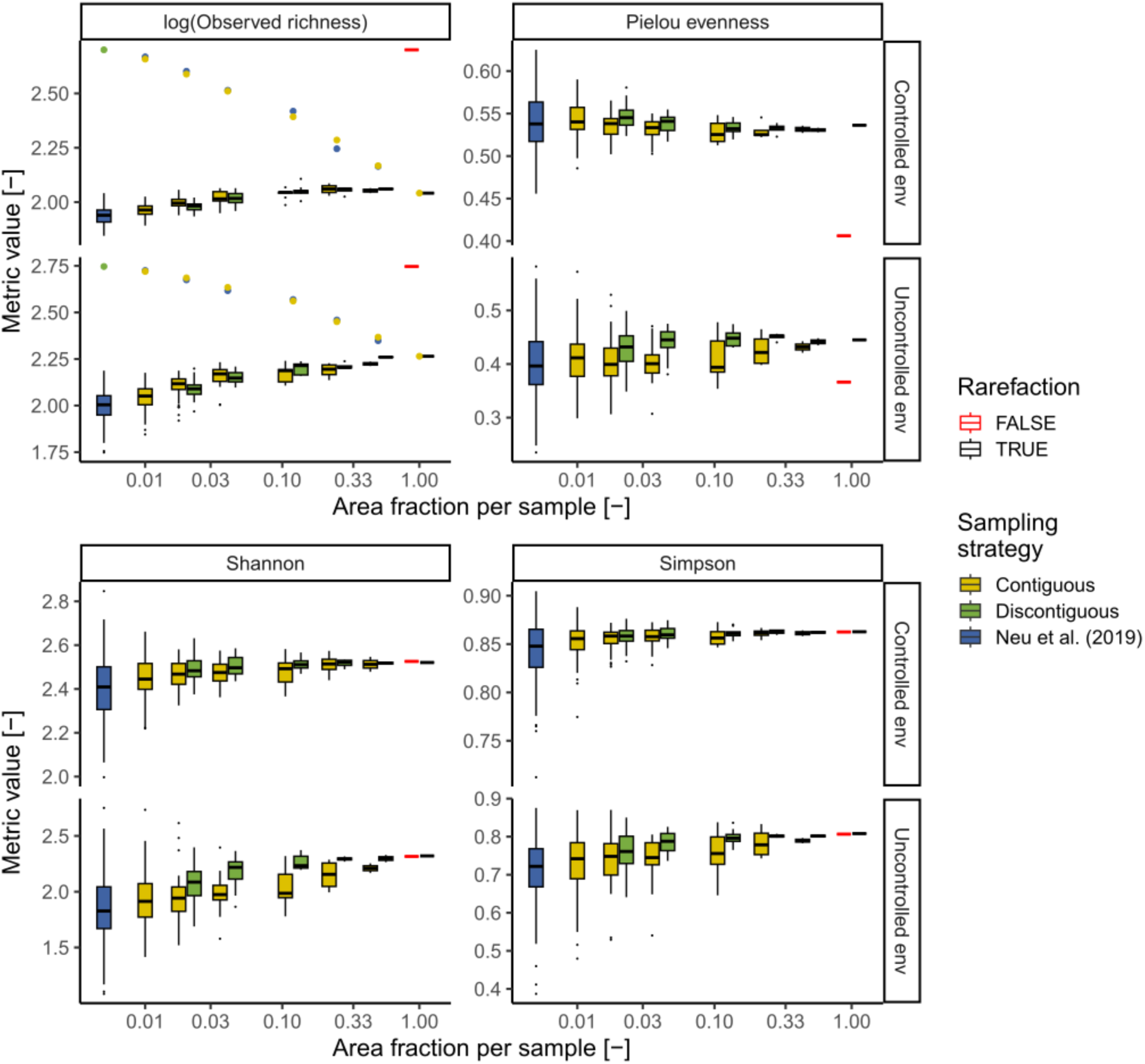
Effect of different sampling strategies on observed biofilm alpha diversity metrics varying the sampling strategy and the fraction of the total area covered by each sample. Boxplots show the distribution of values in each biofilm section, while points indicate the overall values throughout the entire environment.

Similarly, while not significantly affecting community composition (PERMANOVA; p-vals > 0.8), we also noticed how increasing sample area leads to reduced heterogeneity in community composition (Test of multivariate homogeneity; p-vals < 0.01) regardless of the sampling strategy used (Figure 5A). In both environments, however, increasing the sample area using random biofilm sections, as carried out by the discontiguous strategy, reduced drastically the community diversity among samples compared to contiguous samples (Figure 5B), reaching values comparable to the controlled environment and indicating an effective disruption of the biogeographic patterns.

**Figure 5.**
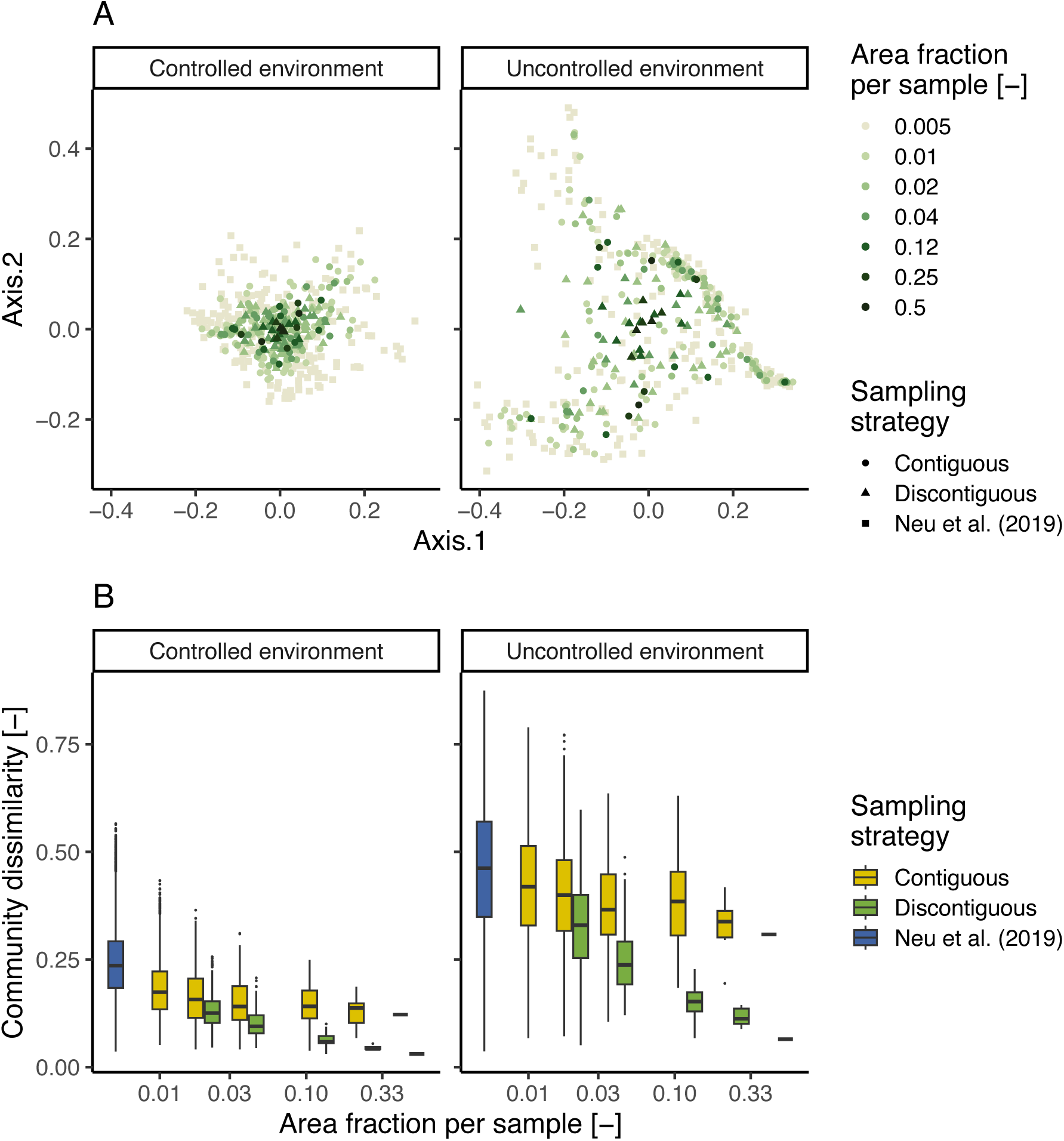
Effect of different sampling strategies on observed biofilm beta diversity metrics. (A) Non-metric multidimensional scaling (NMDS) projection of the Bray-curtis diversity among samples varying the area covered by each sample and sampling strategy. (B) Distribution of beta diversity values varying the area covered by each sample and sampling strategy.

Besides diversity estimates, sampling larger biofilm areas also affected the estimated occupancy of different taxa and their co-occurrence (Figure 6). In fact, we found that sampling larger contiguous biofilm areas leads to inflating the number of co-occurring zOTU pairs, as well as the number of zOTUs which are found ubiquitously in an environment. This result arises due to the spatial pooling performed when sampling larger areas which leads to a simplification of biofilms’ spatial structure. The disruption of the biogeographic patterns obtained by the discontiguous sampling strategy resulted in a drastic increase in zOTUs detected in all samples. Given that the co-occurrence method chosen (i.e., CooccurrenceAffinity) does not consider in its calculations taxa which are present in all samples, this reduced the percentage of co-occurring zOTUs compared to the contiguous sampling strategy. Yet, increased discontiguous sample areas still resulted in an artificial increase of co-occurring zOTUs. Compared to the number of co-occurring taxa, the percentage of zOTUs increased less dramatically with increasing areas per sample due to the reduced sequencing depth over the overall environments and its effect on the number of detected taxa (Figure 4).

**Figure 6.**
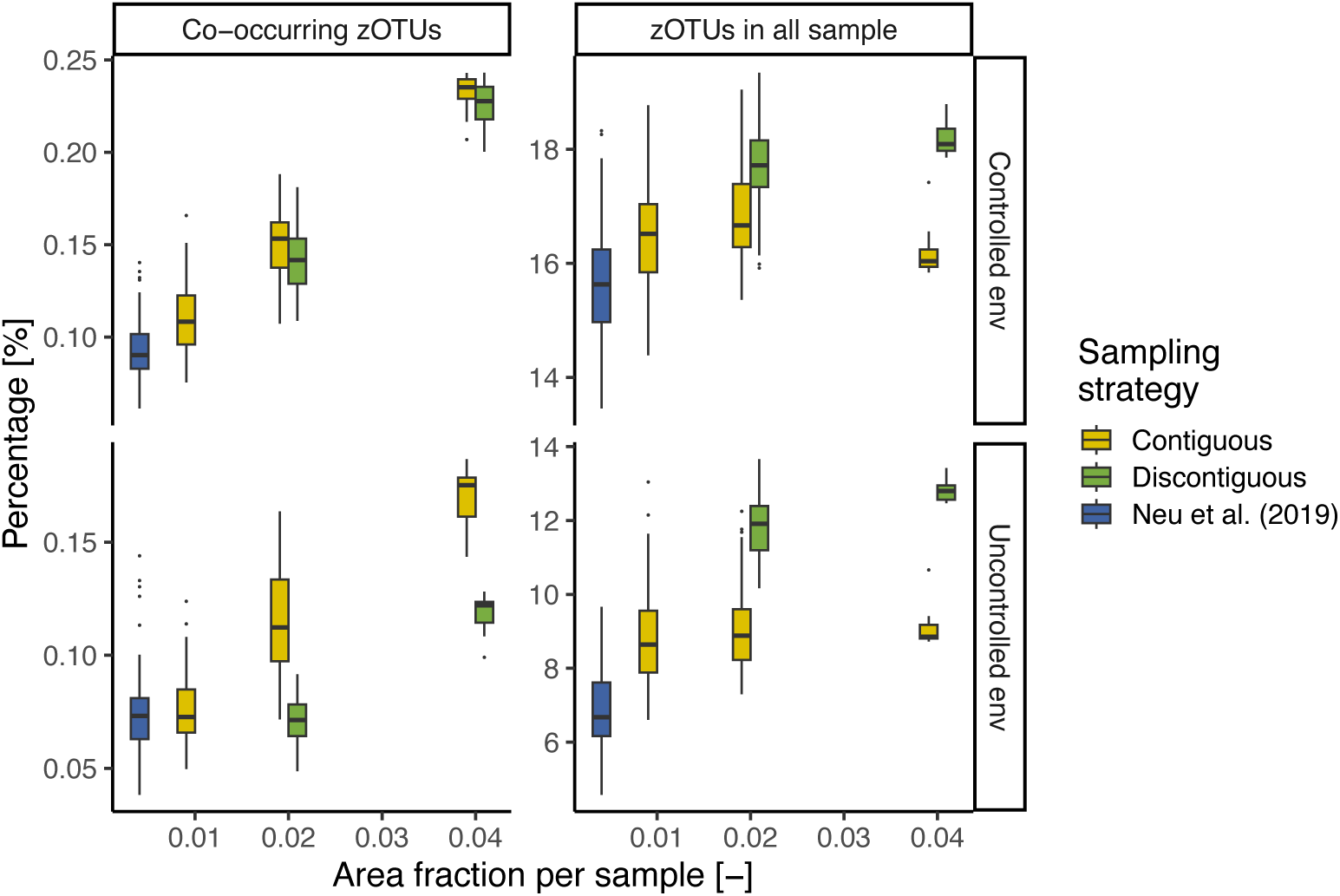
Number of statistically co-occurring zOTUs pairs and zOTUs present in all samples in the controlled and uncontrolled environment as a function of the sampling strategy and the area of biofilm per sample.

Noteworthy, the decrease in heterogeneity at larger sample area was still observed in case no rarefaction was performed, indicating that this phenomenon is not due purely to the decrease in sequencing depth throughout an environment, but rather the spatial averaging of heterogeneous sections (Figure S5).

### The applicability of scaling laws is influenced by environment type and sampling scheme

The increase of taxonomic richness at increasing areas is a well-accepted concept within ecology [45]. However, several different scaling laws have been proposed to describe such species-area relationships (SAR) [40]. Given the cm-scale spatial resolution at which the microbial community composition was collected (Figure 1), this dataset provides an ideal opportunity to test the applicability of different scaling laws to biofilm developed in different types of environments and the impact of different sampling strategies on SAR patterns.

When testing two different scaling laws, namely the power law (PL) and the logarithmic power law (LPL), we found that their fitting error depended on the type of environment (Figure 7A). The power law exhibited the lowest root mean square error (RMSE) in the controlled environment (RMSEPL = 2.4, RMSELPL = 23.9), while the logarithmic power law performed best in the uncontrolled environment (RMSEPL = 13.2, RMSELPL = 2.9). This difference was confirmed when simulating synthetic taxa distributions based on the two environments (Figure S6; Wilcox test; p-vals < 0.001). The difference between the two scaling laws is that the logarithmic power law predicts a faster richness accrual than the power law. A faster accrual of richness is expectable at in an environment with greater variability in environmental conditions and niche differences, leading to a lower overlap between the number of species in different areas than in an homogeneous environment. Such a result is concordant with the greater heterogeneity among microbial communities observed in the uncontrolled environment (Figure 2). We found that, when using the best-performing scaling law for each environment, observing less than 10% of the area was sufficient to estimate the total observed richness with an error below 10%, highlighting the importance of selecting the appropriate scaling law when fitting SAR curves.

**Figure 7.**
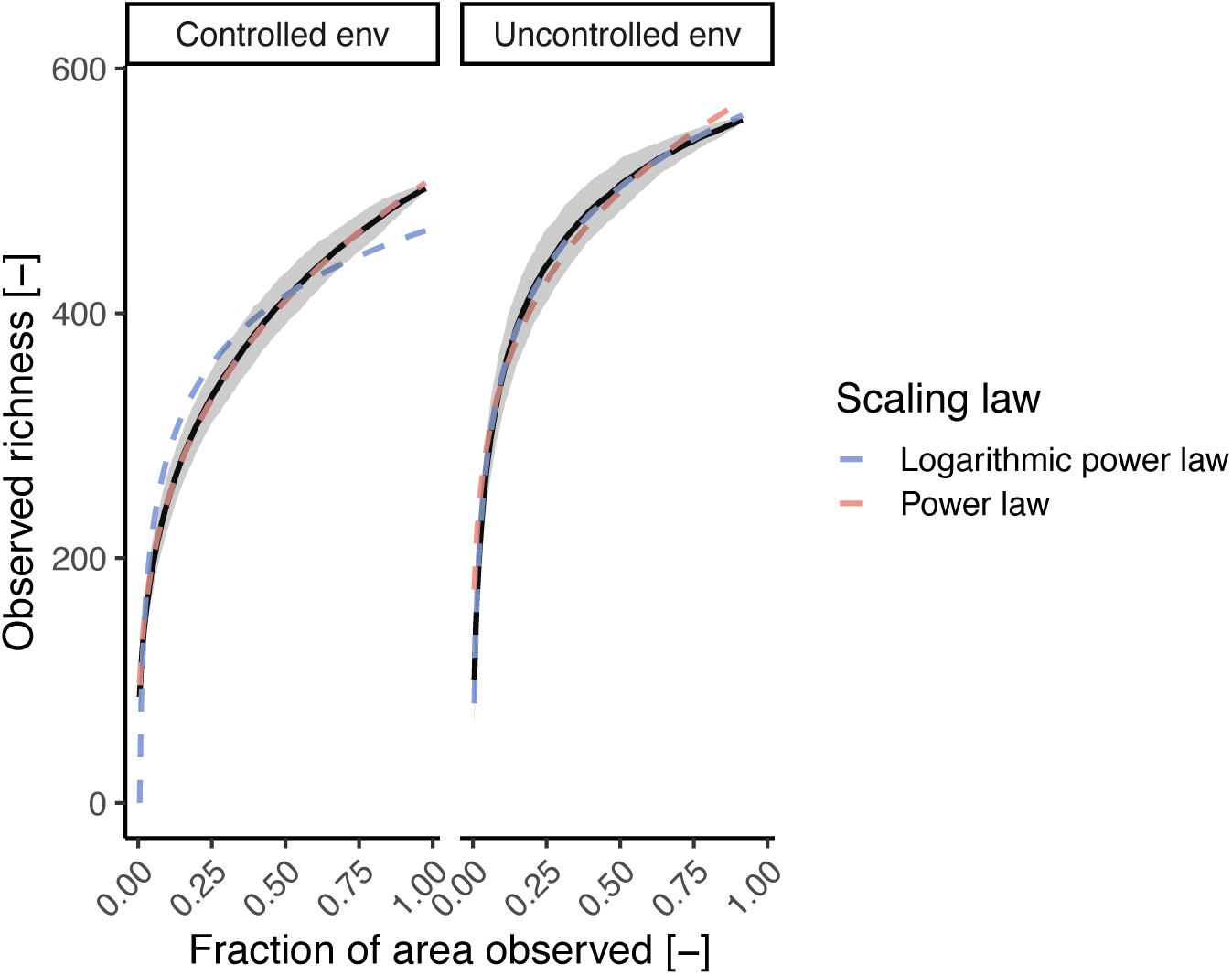
Observed zOTU richness and its standard deviation as a function of the fraction of area observed in each environment (black line and grey area) and predicted species accumulation curves fitted using the power law (PL) and the logarithmic power law (LPL).

Besides the type of environment, the sample size affected the accuracy of scaling law fitting and the estimate of the parameter *z* (Eq. 2) which represents the spatial turnover (i.e., the replacement of taxa from one location to another) (Figure 7). In the controlled environment, we found the RMSE of the power law to be close to zero and smaller than the one provided by the logarithmic power law regardless of the area per sample used (Figure 7B). In the uncontrolled environment, larger sample areas showed a decrease of the power law RMSE and an increase of the logarithmic power law RMSE, resulting in a comparable RMSE at the larger sample area size. Such a different behavior is likely caused by the averaging effect of larger sample areas on the characteristics of local microbial communities in uncontrolled environments. Conversely, despite the presence of biogeographic patterns (Figure 2D), the community structure of the controlled environment is more homogeneous than the uncontrolled one, leading, thus, to observation being less affected by sample size. This is further exemplified by the difference of RMSE values in the uncontrolled environment obtained by the discontiguous sampling strategy at larger areas per sample. Since this sampling strategy pools together randomly-selected biofilm sections disrupting biogeographic patterns, the samples approach diversities comparable to a macroscopically-homogeneous environment (Figure 5B) resulting in a better fit of the power law compared to the logarithmic power law.

Besides scaling law fit, the size of each sample affected the estimated spatial turnover rates. This was especially relevant in the case of the logarithmic power law, where the values varied up to 187% (Figure 7C). The spatial turnover estimated by the logarithmic power law exhibited a maximum at medium sample sizes (i.e., 2-5% of the whole area) and then decreased in both environments. While the diverse sampling strategies affected the predicted spatial turnover, the observed trend did not vary drastically among them. Conversely, the estimates for the power law, while varying less with sample size, showed different trends in the two environments depending on the sampling strategy used. In the controlled environment, regardless of the sampling strategy, the estimated spatial turnover increased with larger sample areas. On the contrary, the contiguous sampling strategy reported a slight turnover decrease at larger areas, while the discontiguous strategy, given its disruption of biogeographic patterns, resulted in a pattern comparable to the controlled environment. The differences observed among sampling strategies and sample areas indicate the importance of taking into account these factors when comparing spatial turnover rates among different studies.

## Discussion

Most environmental microorganisms reside in biofilms [1] and understanding the ecology of these systems is key for both fundamental and applied microbiology. In our study, we show heterogeneity in microbial community composition within meter-scale connected environments of potable water pipes, affected by the variability in localized environmental conditions. While previous reports highlighted how physical separation (e.g., islands) are required to stimulate microbial diversity (e.g., [46]), our results show that, despite being continuously connected, biofilms are composed of heterogeneous neighborhoods of limited diversity (Figures 2-3). Our diversity estimates in local biofilm neighborhoods (Figure 3) align with multiple studies highlighting low local diversity within biofilms [12,13]. Such a low diversity limits the interactions of single cells and their progeny with other species and is likely partially responsible for the non-randomness in the relative abundance pattern. While the location at which a microorganism attach to a biofilm is governed mostly by flow conditions and likely random, establishment (and thus growth) is likely to be dictated by the microorganism’s response to its local neighborhood. For example, taxa which depend on nutrients provided by other taxa, such as auxotrophs, will establish only in neighborhoods were such nutrient-sharing taxa are present [43]. As a result of such these interactions, continuous immigration, while potentially reducing the biogeographic pattern of a biofilm due to the randomness of immigrants’ attachment, cannot completely remove biogeographic patterns arising from the selective effect of biotic interactions on taxa establishment. Hence, biofilms that are mostly sustained by growth and experience limited dispersal (i.e., continuous immigration and emigration) will likely present greater heterogeneities, resulting in patches likely dominate by few species.

As a result of the spatial heterogeneity of biofilm communities, the sample area has profound implications for the study of biofilms: sequencing results from a sample report only its “averaged” composition and cancel out local heterogeneities and associations when the sample area is too large, or miss key communities when too small. As a result, a large sample size led to an overestimation of the co-existing taxa and an inflated number of statistical associations (Figure 6). For example, while multiple related zOTUs have been reported to colonize real shower hoses biofilms when sampled as a whole [47], the increased spatial resolution of this study highlighted how related taxa tend to occur in different biofilm locations (Figure S3), likely depending on their distinct environmental preferences and interactions. This underlines the requirement for a biologically-based definition of sampling units. For example, while associations networks have often been used to infer interactions in biofilms (e.g.,[47,48]), such practices should rely on sample sizes informed by the spatial limits of most microbial interactions [9], as for example achievable using SAMPL-seq [49]. Moreover, the impact of sample sizes on community characteristics and estimated co-occurrence (Figure 4-6), as well as the estimated spatial turnover (Figure 8) calls for the use of standardized sampling areas within a study and cautions against cross-study comparisons when sample sizes differed.

**Figure 8.**
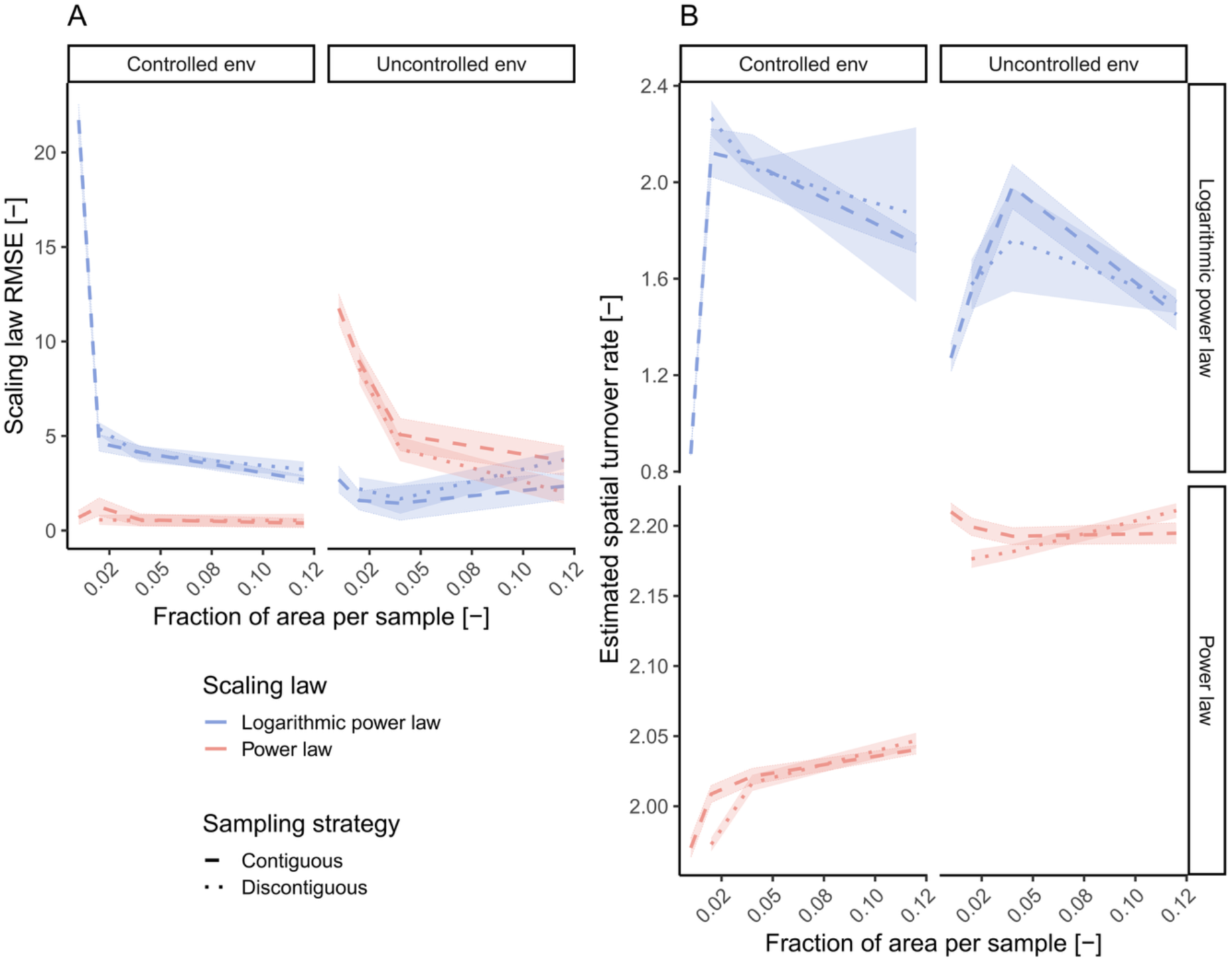
(A) Scaling law RMSE of the sampling schemes tested as a function of the fraction of area observed in each environment. (B) Estimated spatial turnover values in each environment as a function of the scaling law and sampling scheme used.

Samples should, ideally, be a representative fraction of the environment of interest from which properties of the whole environment could be estimated. The presence of biogeography in biofilms not only hinders the identification of a correct spatial scale [11], but also the selection of a representative fraction. For example, even if the fluctuation of community structure in the overall similar control environment could be considered as negligible, the Bray-curtis diversity observed in the satellite community composition reach values above 0.75 (Figure 2), indicating dramatically different microbial communities at 1.2 m distances. One approach could be to enlarge the area per sample, especially in environments which could be sampled entirely (e.g., shower hoses). An alternative strategy would be to circumvent the problem of identifying a single “representative sample” per environment and rely on few samples of small areas distributed uniformly within the environment of interest. Given the biogeography within biofilms, such samples should not be considered as biological replicates, but rather as a measure of the biofilm heterogeneity within a given environment. Such samples could either be pooled to obtain a single sample disrupting biogeographic patterns and reflecting an “averaged” microbial communities within an environment or, better, included independently in downstream statistical analyses, allowing to take explicitly into account communities heterogeneity within an environment. All sampling strategies are characterized by pros and cons which depend on the research question at hand. Enlarging the area per sample (both in a contiguous or discontiguous way) confounds the study of microbial interactions given its averaging effect on community composition. Conversely, when paired with adequate sequencing depth, such a strategy allows to sample more local neighborhoods per sample, increasing the diversity of detected taxa and aiding in the detection of taxa of specific interest (e.g., pathogens). While sampling contiguous areas would be sufficient in environments with low variability, in case of expected changes of environmental conditions it would be more appropriate to distribute randomly the sampling area across the whole environment in order to reduce the effect of biogeography. On the other hand, sampling independently few small biofilm areas distributed across an environment permits to evaluate the heterogeneity within an environment and, limiting spatial averaging, is more suited to study biofilm ecology of local biofilm neighborhoods or upscale biofilm characteristics to the whole environment. On the other hand, in case of pronounced biofilm heterogeneity such a strategy might miss specific taxa of interest present in areas not sampled.

In conclusion, this work demonstrates the presence of small-scale biogeography in both macroscopically homogeneous and heterogeneous environments, as well as their implications in microbial analyses. Our results call for researchers, even when working in small and seemingly homogeneous environments, to not ignore biofilm heterogeneities and adapt their sampling strategy based on the specific research question at hand.

## Acknowledgements

We recognize the authors of the original dataset for publicly sharing their results. We would also like to thank the members of the Drinking Water Microbiology group and Environmental Microbiology Department at Eawag for the thoughtful discussions on the manuscript, and Eawag discretionary funding for financial support.

## Data availability

Sequencing data from Neu et al. [18] is deposited on NCBI under PRJNA554997, while the scripts for the analyses performed are available at https://github.com/mgabriell1/SmallScaleBiogeography_Biofilms_manuscript_scripts.

