## Supplementary information for "Small-scale biogeography of biofilms and its implications for sequencing-based studies"

* Corresponding author:

Name: Marco Gabrielli

Table S1. Number and percentages of zOTUs with show non-random (i.e., peaking at specific locations) or monotonic (i.e., increasing/decreasing) patters over at least one side of the hoses.

| **Uncontrolled environment** | | Trend | |  | **Controlled environment** | | Trend | |
| --- | --- | --- | --- | --- | --- | --- | --- | --- |
|  |  | TRUE | FALSE |  |  |  | TRUE | FALSE |
| Non-random pattern | TRUE | 69 (12%) | 57 (10%) |  | Non-random pattern | TRUE | 38 (8%) | 30 (6%) |
|  | FALSE | 22 (4%) | 410 (74%) |  |  | FALSE | 31 (6%) | 403 (80%) |

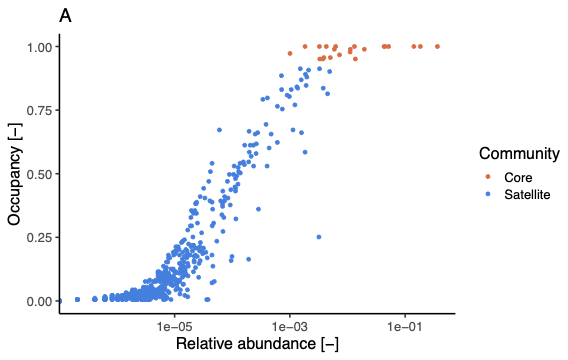

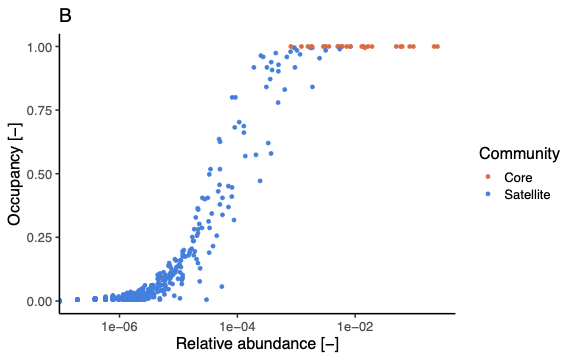

Figure S1. Occupancy and relative abundance of zOTUs in uncontrolled (A) and controlled (B) environment split by core and saltellite communities.

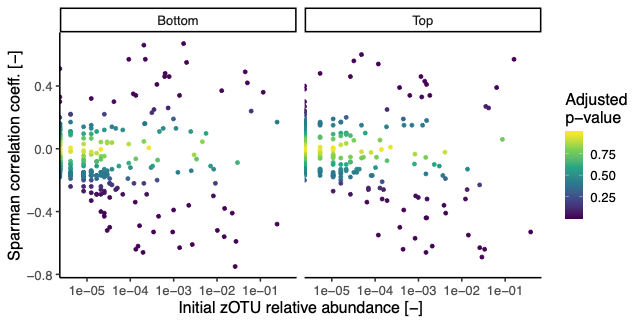

Figure S2. Correlation between zOTUs relative abundance and position along the biofilm as a function of the zOTUs relative abundance over the initial 12 cm of biofilm.

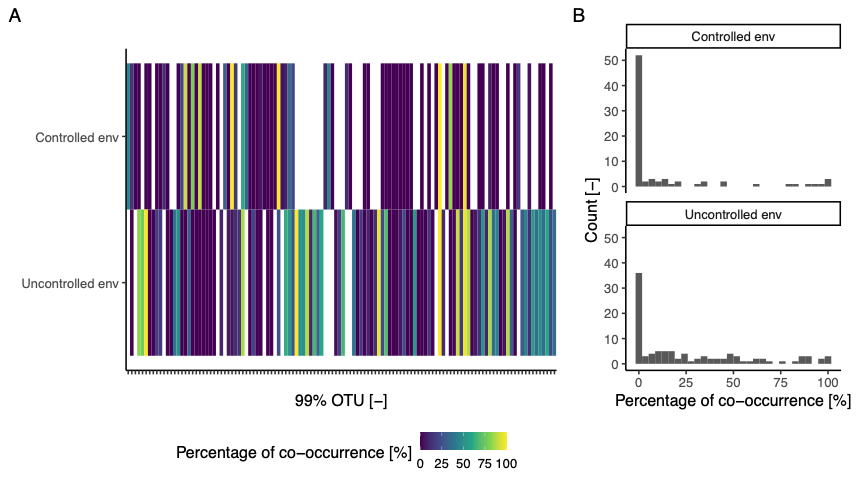

Figure S3. (A) Co-occurrence percentage of zOTUs into 99% similar OTUs. White tiles indicate a missing OTU in a given dataset. (B) Histogram showing the percentage of within-OTU co-occurrences in both environments.

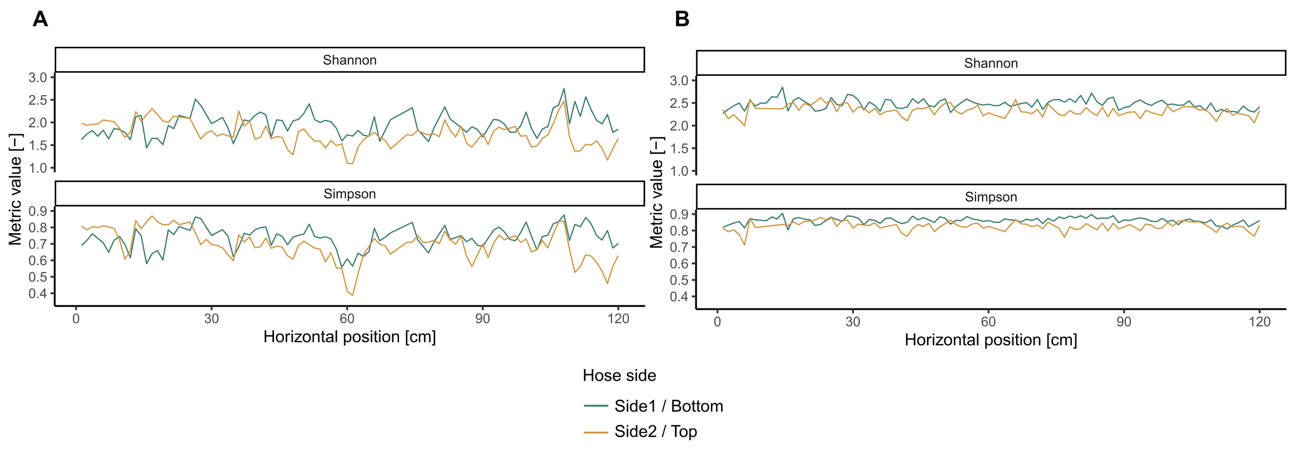

Figure S4. Shannon and Simpson diversity over the biofilms lengths in the uncontrolled (A) and controlled (B) environments.

Table S2. Spearman correlation and p-values (within brackets) along the biofilms’ sides developed in the controlled and uncontrolled environments.

| Metric | Biofilm side | Controlled environment | Uncontrolled environment |
| --- | --- | --- | --- |
| Observed richness | Bottom / Side1 | -0.63 (< 0.001) | 0.28 (0.006) |
|  | Top / Side2 | 0.01 (0.91) | 0.28 (0.006) |
| Pielou’s evenness | Bottom / Side1 | 0.08 (0.45) | 0.16 (0.13) |
|  | Top / Side2 | -0.29 (0.005) | -0.52 (< 0.001) |
| Shannon | Bottom / Side1 | -0.16 (0.10) | 0.26 (0.01) |
|  | Top / Side2 | -0.29 (0.004) | -0.47 (< 0.001) |
| Simpson | Bottom / Side1 | -0.11 (0.29) | -0.43 (< 0.001) |
|  | Top / Side2 | 0.23 (0.02) | -0.03 (0.75) |

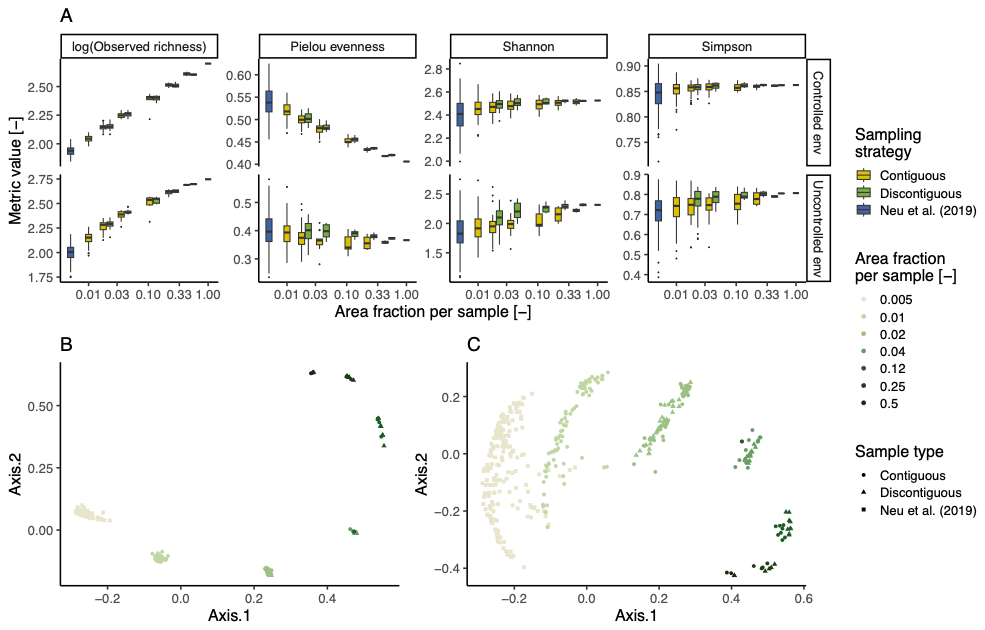

Figure S5. Effect of different sampling schemes on observed biofilm characteristics varying the sampling strategy and the fraction of the total area covered by each sample without considering limitation on sequencing depth per sample. (A) Alpha diversity metrics. Boxplots show the distribution of values in each biofilm section, while points indicate the overall values throughout the entire environment. (B, C) Non-metric multidimensional scaling (NMDS) projection of the Bray-curtis diversity among biofilm sections in the controlled (B) and uncontrolled (C) environments.

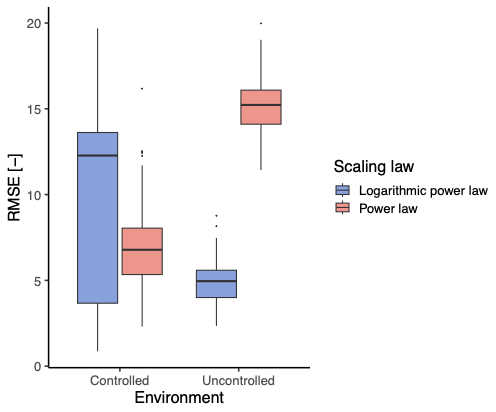

Figure S6. Root mean square error (RMSE) obtained fitting the power law and the logarithmic power law scaling functions on the simulated community composition of biofilm sections obtained using using MIDAsim.
